# A gut microbial metabolite of dietary polyphenols reverses obesity-driven hepatic steatosis

**DOI:** 10.1101/2021.09.16.460661

**Authors:** Lucas J. Osborn, Karlee Schultz, William Massey, Beckey DeLucia, Ibrahim Choucair, Venkateshwari Varadharajan, Kevin Fung, Anthony J. Horak, Danny Orabi, Ina Nemet, Laura E. Nagy, Zeneng Wang, Daniela S. Allende, Naseer Sangwan, Adeline M. Hajjar, Christine McDonald, Philip P. Ahern, Stanley L. Hazen, J. Mark Brown, Jan Claesen

## Abstract

The molecular mechanisms by which dietary fruits and vegetables confer cardiometabolic benefits remain poorly understood. Historically, these beneficial properties have been attributed to the antioxidant activity of flavonoids. Here, we reveal that the host metabolic benefits associated with flavonoid consumption actually hinge on gut microbial metabolism. We show that a single gut microbial flavonoid catabolite is sufficient to reduce diet-induced cardiometabolic disease burden in mice. Dietary supplementation with elderberry extract attenuated obesity and continuous delivery of the catabolite 4-hydroxphenylacetic acid was sufficient to reverse hepatic steatosis. Analysis of human gut metagenomes revealed that under one percent contains a flavonol catabolic pathway, underscoring the rarity of this process. Our study will impact the design of dietary and probiotic interventions to complement traditional cardiometabolic treatment strategies.

**One-Sentence Summary:** Select gut microbes can metabolize flavonoids from a fruit and vegetable diet to monophenolic acids, which improve fatty liver disease.

**Graphical abstract:** 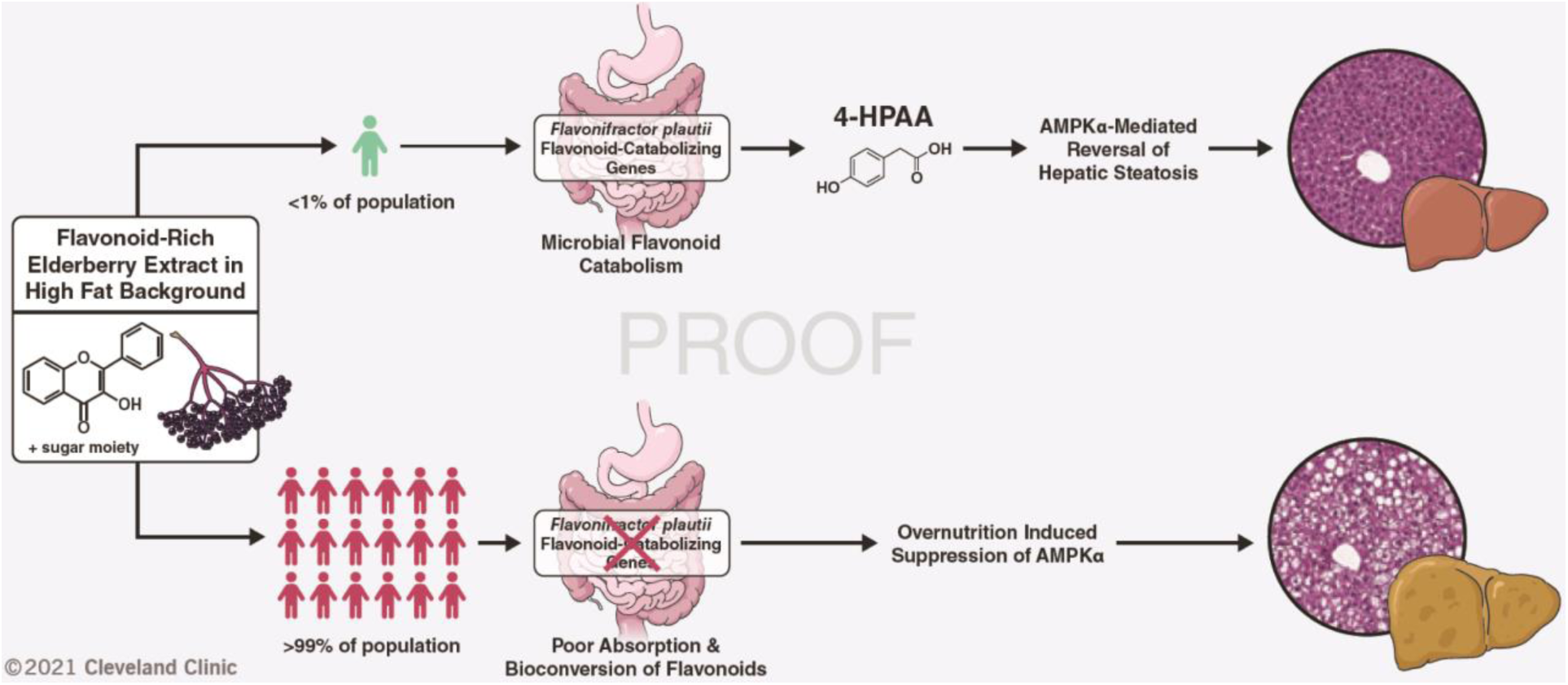

## Main Text

A balanced diet is one of the most influential drivers of human health. This notion has become increasingly important in our industrialized era, characterized by pervasive human metabolic syndrome. More recently, diet has become appreciated as a focal determinant of gut microbial community structure, function, and resilience where dietary choices are recognized to rapidly alter the human gut microbiome (*1*). Moreover, diet-derived metabolites from the human gut microbiota have been causally linked to cardiovascular and metabolic disease pathogenesis (*2*– *5*). Consequently, microbial metabolites arising from specific dietary components, such as trimethylamine (TMA), imidazole propionate and short-chain fatty acids (SCFAs), have gained recognition as central mediators of human health and disease (*2–6*).

Flavonoids represent a key molecular component of plant-based diets. They have been attributed antioxidant, antiobesogenic, and chemoprotective properties through the scavenging of free radicals and activation of molecular effectors implicated in human disease (*7*). Dietary flavonoids are largely glycosylated, limiting their absorption in the small intestine and thereby their systemic distribution (*7*). Consequently, upon passing into the colon, flavonoids become a substrate for gut microbial catabolism. Notable prior studies have highlighted that dietary flavonoids attenuate diet-induced obesity in a microbe-dependent manner (*8, 9*). Several human gut bacteria that are capable of breaking down flavonoid substrates through reduction and subsequent cleavage of the central non-aromatic ring, followed by hydrolysis to yield monophenolic acid degradation products have been isolated (*10*). The four types of bacterial enzymes required in the flavone/flavonol catabolic pathway (flavone reductase (FLR), chalcone isomerase (CHI), enoate reductase (EnoR), and phloretin hydrolase (PHY)) have been identified and biochemically characterized (*11–14*). Together, these genes represent just one method by which commensal gut bacteria metabolize host dietary inputs; additional homologous or non-homologous pathways may remain undiscovered.

### Berry supplementation reduces the obesogenic effects of a high fat diet

Since ingested flavonoids themselves have poor bioavailability, we hypothesized that their microbial monophenolic acid catabolites are responsible for prior ascribed anti-obesogenic properties (*8, 9*). To identify candidate catabolites, we used a comparative targeted metabolomics analysis of mice that were supplemented three different flavonoid-rich berry extracts on a high fat diet (HFD) background. For 16 weeks, we provided mice with ad libitum access to a high fat control diet, or the same high fat diet base, supplemented with 1% w/w of either elderberry, black currant, or aroniaberry extracts. Metabolic parameters including body weight, lean mass, fat mass, and glucose homeostasis were tracked for the duration of the experiment. We observed that elderberry extract-supplemented mice were markedly protected from HFD-induced obesity (Fig. 1A-D, S1A, B). Moreover, elderberry extract-supplemented mice had more lean mass, less fat mass, and were less hyperinsulinemic than HFD control mice 16 weeks after initiation of differential diet treatments (Fig. 1E-G, S1C). Analysis of the cecal microbial composition by 16S rRNA sequencing revealed that elderberry extract-supplemented mice had significantly more diverse cecal microbial communities that clustered distinctly from the HFD control mice using non-metric multidimensional scaling, and from the other berry extract-supplemented mice (Fig. 2A-C, S2A-I). Of note, *Lachnospiraceae UCG-006* was only observed in the cecal communities of berry-extract fed mice (light green in Fig. 2C, S2F). Moreover, other *Lachnospiraceae* clades (*NK4A136* and *UCG-009*) were significantly more abundant in elderberry-extract fed mice when compared to the HFD control group (Fig. 2C, S2G).

**Fig. 1.**
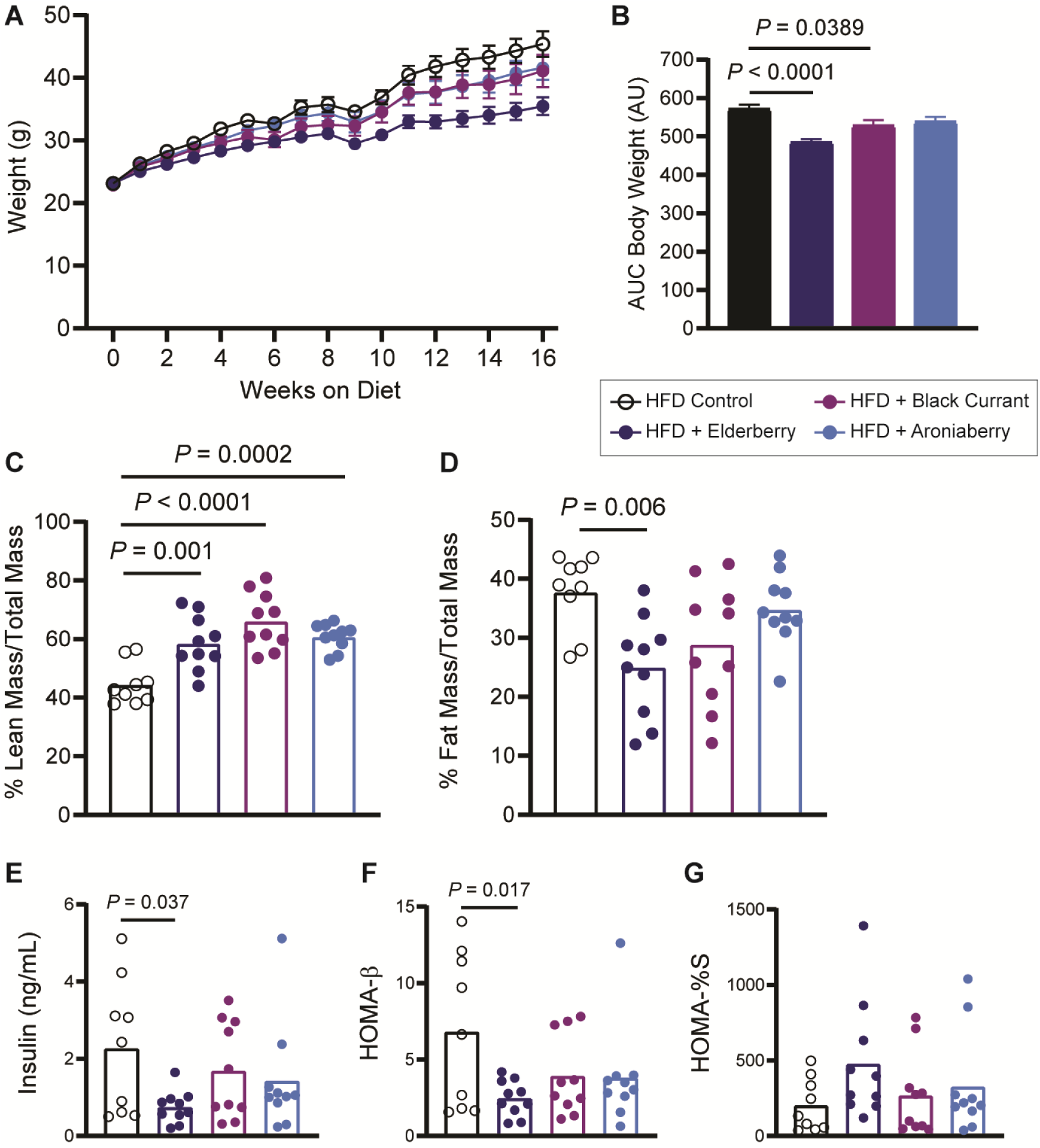
Berry extract supplements reduce high fat diet-induced obesity. **A)** Body weights of 6-week-old male C57BL/6 mice fed a high fat control diet, or the same diet supplemented with 1% w/w elderberry, black currant, or aroniaberry extracts for 16 weeks; n=9-10 per group, error bars represent SEM. (**B**) Mean cumulative area under the curve (AUC) for body weights after 16 weeks, error bars represent SEM. (**C**) **and** (**D**) Lean and fat mass after 16 weeks as measured by EchoMRI and normalized to total body mass, n=9-10 per group. (**E**) Plasma insulin after 16 weeks on either control or experimental diets following a 4-hour fast, n=9-10 per group. (**F**) Homeostatic model assessment of β-cell function (HOMA-β), n=9-10 per group. (**G**) Percent insulin sensitivity after 16 weeks, n=9-10 per group. Individual points represent individual mice, and bars represent group means. All *P* values shown were calculated using one-way ANOVA with Dunnett’s multiple comparisons test; n=9-10 per group.

**Fig. 2.**
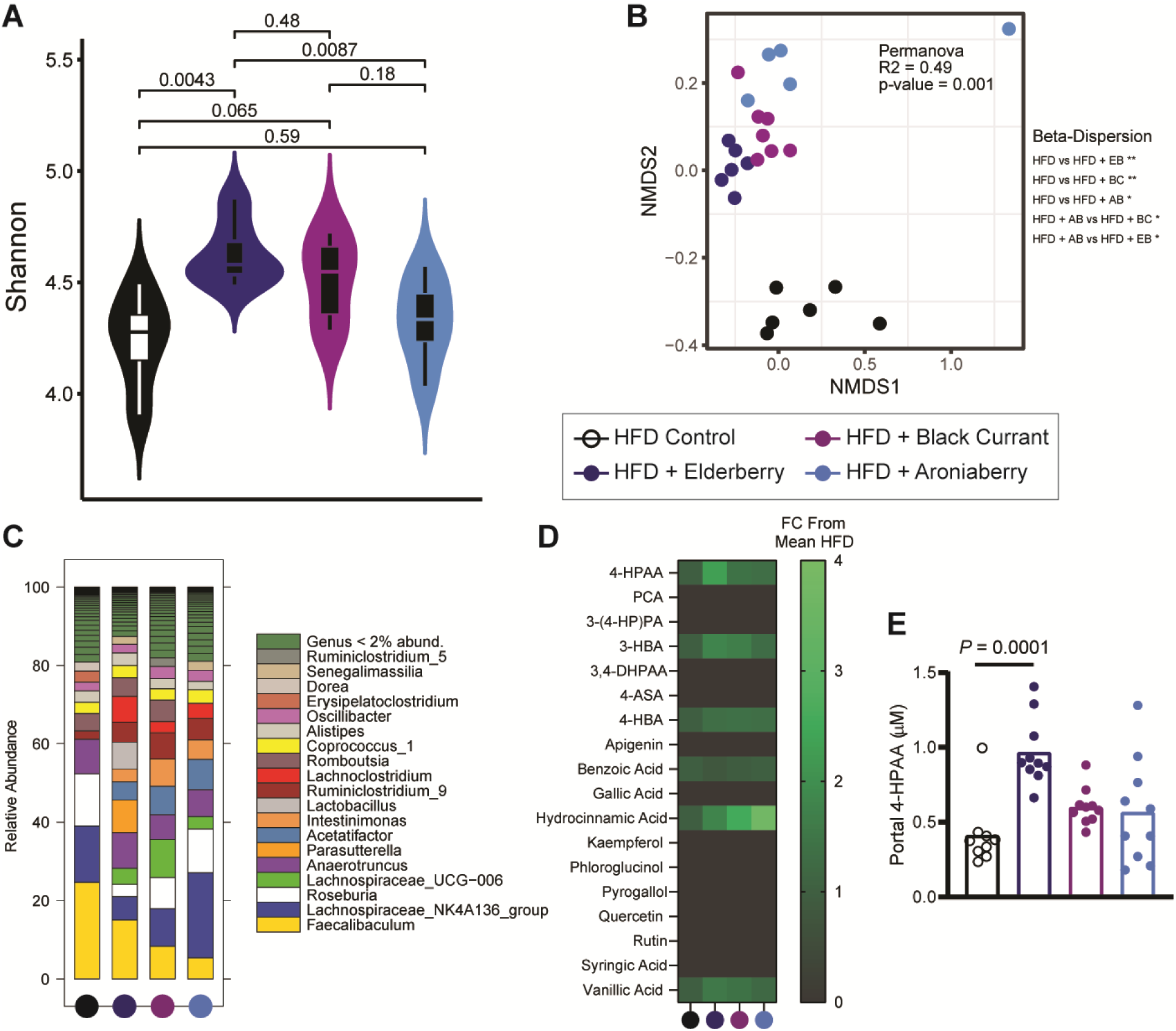
Berry extracts alter the cecal microbiome. (**A**) Shannon alpha diversity estimates for cecal microbiomes based on 16S rRNA profiles of all four groups. Statistical analysis was performed via ANOVA. (**B**) NMDS plots based on the Bray-Curtis index between the cecal 16S rRNA profile of all four groups. Statistical analysis was performed with PERMANOVA where R2 values are noted for comparisons with significant p-values and stand for percentage variance explained by the variable of interest. (**C**) Stacked bar chart of relative abundance (left y-axis) of the top 20 genera assembled across the cecal 16S rRNA profiles of all four groups. n=6 for all 16S rRNA sequencing analysis. (**D**) Heatmap of portal plasma flavonoids and microbial flavonoid catabolites measured by LC-MS/MS, n=9-10 per group. (**E**) Portal plasma concentration of the microbial flavonol catabolite 4-HPAA measured by LC-MS/MS, one-way ANOVA with Dunnett’s multiple comparisons test; n=9-10 per group. Individual points represent individual mice, and bars represent group means.

Provided that mice are likely to contain commensal gut microbes capable of catabolizing flavonoids into monophenolic acids, we expected that these microbe-derived monophenolic acids would be enriched in the portal plasma of berry extract-supplemented mice compared to the HFD control. A targeted liquid chromatography-tandem mass spectrometry (LC-MS/MS) approach revealed six known microbial flavonoid catabolites at detectable levels in portal plasma (*15*) (Fig. 2D, S2J-N). Of the six metabolites, only 4-hydroxyphenylacetic acid (4-HPAA) and 4-hydroxy-3-methoxybenzoic acid were significantly enriched in the portal plasma of elderberry-supplemented mice compared to the HFD control (Fig. 2D, E, S2J). Between these two metabolites, 4-HPAA had the largest fold difference in portal plasma concentration relative to the HFD control (2.33 +/- 0.35, mean fold change +/- 95% confidence interval, n=9-10). In addition, 4-HPAA correlated negatively with plasma insulin levels (Fig. S1C) (R^2^=0.206, *P*=0.004). Finally, plasma levels of 4-HPAA have been previously reported to associate negatively with indices of obesity in a cohort of non-diabetic obese human subjects (*16*). Hence, we selected 4-HPAA as our candidate molecule to test the hypothesis that a single microbial flavonoid catabolite would be sufficient to abrogate key parameters of HFD-induced metabolic disease.

### A single microbial flavonoid catabolite reverses hepatic steatosis

To test whether the metabolically beneficial effects of elderberry supplementation could at least in part be attributed to the microbial flavonoid catabolite 4-HPAA, we implanted subcutaneous slow-release pellets delivering 4-HPAA (350 μg/day) into mice pre-fed a high fat diet to establish obesity and compared them to obese control mice that received an implanted sham scaffold for 6 weeks (Fig. 3A). This subcutaneous delivery method has been used previously to study different microbial metabolites at a single-metabolite resolution (*3*). Its main advantage is that the delivered molecule is independent of further modification by gut microbial metabolism, a limitation of providing 4-HPAA in the diet or drinking water (*3*). After twenty-five days, we surveyed global metabolism and energy substrate utilization using indirect calorimetry data to compare the 4-HPAA and the scaffold-only control mice at isothermal 30 °C, room temperature 23 °C, and under cold challenge at 4 °C. We observed that 4-HPAA treated mice were more prone to utilize carbohydrates as an energy source as measured by the respiratory exchange ratio (RER) during the light cycle under cold exposure (Fig. S3A). At the time of sacrifice, the mRNA expression of *Ucp1* (encoding a key regulator of non-shivering thermogenesis that mediates oxygen consumption in cold conditions (*17*)) was increased in the metabolically active brown adipose tissue of 4-HPAA treated mice (Fig. S2B). This suggests that 4-HPAA may modulate metabolic flexibility during cold exposure.

**Fig. 3.**
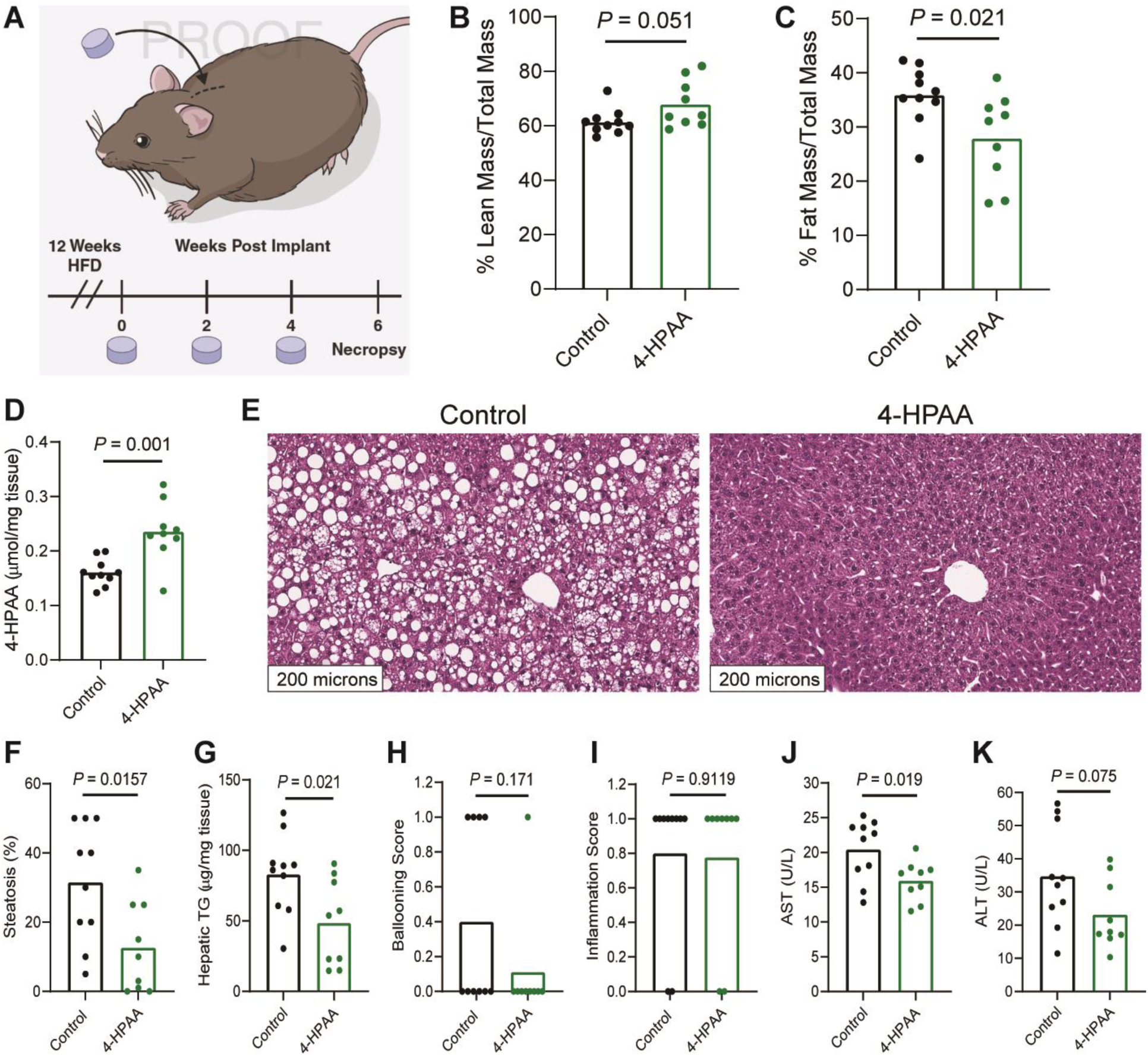
4-HPAA is sufficient to reverse HFD-induced hepatic steatosis and liver injury. (**A**) 4-week-old male C57BL/6 mice were fed a high fat diet for 12 weeks to induce obesity. After the induction of diet-induced obesity, mice were randomly assigned to receive a subcutaneously implanted control scaffold pellet, or a pellet releasing 350 μg 4-HPAA per day for two weeks. New pellets were implanted every two weeks for a total of 6 weeks. Lean mass (**B**) and fat mass (**C**) were measured by Echo MRI, n=9-10 per group. (**D**) 4-HPAA accumulates in the liver as measured by LC-MS/MS, n=9-10 per group. (**E-G**) After a total of 18 weeks of high fat diet feeding, H&E-stained sections of liver revealed profound hepatic steatosis in scaffold control mice while 4-HPAA treated mice had marked reversal of steatosis and triglyceride deposition n=6-10 per group. Histologic assessment of hepatocellular ballooning (**H**) and inflammation (**I**), n=6 per group. (**J, K**) Quantification of liver injury biomarkers in peripheral plasma, n=9-10 per group. Statistical analysis was performed using unpaired two-tailed Student’s t-test. Individual points represent individual mice, and bars represent group means.

After six weeks of continuous 4-HPAA exposure, modest changes in gross anthropometrics (Fig. 3B, C) were overshadowed by striking liver-specific effects of 4-HPAA. Consistent with our previous findings (*18*), 4-HPAA accumulates in the liver as it undergoes rapid first-pass hepatic metabolism (Fig. 3D). Remarkably, after just 6 weeks, subcutaneous 4-HPAA administration led to a marked reversal of hepatic steatosis when compared to control mice (Fig. 3E-G). We did not observe quantitative differences in the mild presentation of hepatocellular ballooning or lobular inflammation (Fig. 3H, I), yet 4-HPAA treated mice had lower plasma concentrations of the liver injury marker aspartate aminotransferase but not alanine aminotransferase (Fig. 3J, K).

To better understand the transcriptional foundations of the 4-HPAA mediated reduction of hepatic steatosis, we measured the mRNA expression of several genes implicated in lipid metabolism and inflammation in the liver (Fig. S3C). Here we observed a reduction in the fatty acid importer *Cd36*, the fatty acid desaturase *Scd1*, and the pro-inflammatory cytokine *Tnfa* in 4-HPAA treated mice with a concomitant increase in the master regulator of mitochondrial biogenesis, *Pgc1a* and *Lcn13*, a regulator of glucose and lipid metabolism (*19–21*). In addition, we measured the mRNA expression of lipid metabolism and inflammation-related genes in the gonadal white adipose tissue as it plays a central role in maintaining metabolic homeostasis (Fig. S3D). For 4-HPAA treated mice, we observed increased mRNA levels of lipoprotein lipase (*Lpl*), the adipocyte-specific isoform of peroxisome proliferator-activated receptor gamma (*Pparg2*, a master regulator of glucose homeostasis and lipid metabolism), as well as a reduction of the expression of macrophage-associated glycoprotein *Cd68*. Taken together, our data suggest that 4-HPAA confers a distinct metabolic benefit that is regulated in part by tissue-specific transcriptional reprogramming.

### 4-HPAA activates hepatic AMPKα signaling

We next set out to uncover the signaling cascade(s) that could be potentially linked to 4-HPAA induced transcriptional changes. We hypothesized that AMP-activated protein kinase (AMPK) signaling is upregulated in the livers of mice receiving subcutaneous 4-HPAA. Our reasoning was based on four premises (i) literature reports on the activation of AMPK by the structurally distinct monophenolic acids gallic acid, vanillic acid and caffeic acid (*22–24*), (ii) the observation that exogenously administered 4-HPAA accumulates in the liver, (iii) our observed transcriptional data that is consistent with AMPK activation (*25*), and (iv) physiologic hormone-driven activation of AMPK by adiponectin and leptin is potently anti-steatotic (*26–28*). We next tested our hypothesis experimentally by immunoblot analysis of the liver tissues from our subcutaneous delivery experiment. This analysis revealed increased phosphorylation of AMPKα (Fig. 4A) and its downstream effector acetyl-CoA carboxylase (ACC, a central mediator of de novo fatty acid synthesis; Fig 4B) in 4-HPAA treated mice when compared to the scaffold control group. This observation was specific to the α subunit of AMPK, as subcutaneous 4-HPAA administration did not lead to the phosphorylation of the AMPKα subunit (Fig. S4).

**Fig. 4.**
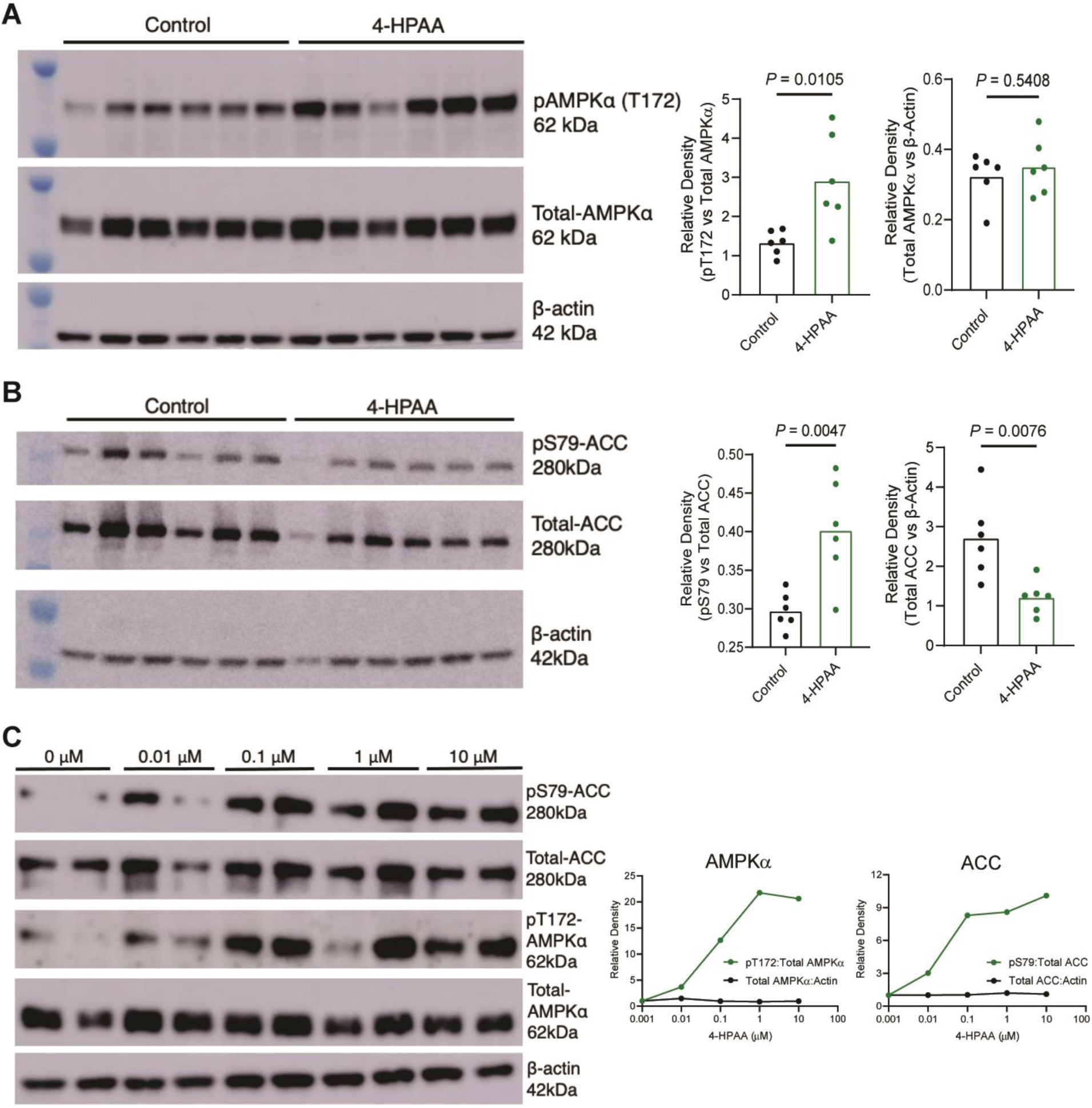
4-HPAA activates AMPKα and downstream effectors to reduce de novo hepatic lipogenesis. (**A)** Western blot analysis of pAMPKα (T172), total AMPKα, and β-actin with densitometric quantification, n=6 per group. (**B**) Western blot analysis of pACC (S79), total ACC, and β-actin with densitometric quantification, n=6 per group. (**C**) Primary murine hepatocytes were treated with either DMSO vehicle control (0 μM) or 0.01, 0.1, 1, and 10 μM 4-HPAA for 30 minutes at which point protein expression of pACC (S79), total ACC, pAMPKα (T172), total AMPKα, and β-actin were measured via Western blot analysis with densitometric quantification with dots representing the mean of two biological replicates. Statistical analysis for panels A and B was performed using unpaired two-tailed Student’s t-test. Individual points represent individual mice, and bars represent group means.

To determine whether 4-HPAA can activate AMPKα in a cell autonomous manner we treated primary mouse hepatocytes with physiologically relevant concentrations of 4-HPAA (0.01-10 μM) for 30 minutes. Mirroring our *in vivo* observations, 4-HPAA treatment of primary mouse hepatocytes lead to the phosphorylation of AMPKα and ACC in a dose-dependent manner *in vitro* (Fig. 4C). Surprisingly, the peak phosphorylation of AMPKα was observed after treatment with 1 μM 4-HPAA, the same concentration as observed in the portal blood of our elderberry extract-supplemented mice (Fig. 4C, 2E). These results indicate that 4-HPAA acts locally in the liver –the primary site of accumulation– to activate AMPKα and downstream signaling events that in turn activates fatty acid oxidation and blunts *de novo* lipogenesis.

### Flavonol catabolism is a rare feature among human microbiota

While our berry-fed mice harbored flavonoid-catabolizing gut microbiota members capable of producing 4-HPAA, clinical studies have shown that flavonoid catabolism is less prominent among human gut microbiota and displays marked inter-individual variation (*29, 30*). Several human gut commensals capable of catabolizing dietary flavonols into monophenolic acids have been identified, the most studied example being the eponymous *Flavonifractor plautii* (formerly *Clostridium oribscindens*) (Fig. S5) (*15, 31*). Notably, *F. plautii* belongs to the *Lachnospiraceae* family, which accounted for one of the major changes in the microbiota of our berry supplemented mice (Fig. 2C). Recently, the complete set of genes required for flavonol catabolic activity was characterized. In total, four genes are required for the stepwise degradation of flavones and flavonols into monophenolic acids such as 4-HPAA. These four genes encode a flavone reductase (FLR), chalcone isomerase (CHI), enoate reductase (EnoR), and phloretin hydrolase (PHY) (*11–14*) (Fig. 5A). To predict the flavonol catabolic capacity of human metagenomes, we calculated the incidence of co-occurrence for this complete catabolic gene set in the metagenomic sequencing data (n=1899 assemblies) from two publicly available repositories (*32, 33*). Strikingly, although *F. plautii* was present in 28 percent of human microbiomes in these datasets (Supplementary Tables 1,2), the incidence of any one catabolic gene (>90% sequence similarity to the reference gene set) does not exceed 3 percent. Moreover, the co-occurrence of all four genes –required for the complete flavonol degradation pathway– is exceptionally rare, with an incidence of roughly one in two hundred (Fig. 5B). These data suggest that not all *F. plautii* strains can catabolize flavonols and support the recent report that *F. plautii* isolates are subject to inter-strain competition in the presence of flavonoids (*34*). Overall, this highlights the rarity of a complete flavonol catabolic gene set in human microbiomes, although the possibility exists that other catabolic pathways remain undiscovered.

**Fig. 5.**
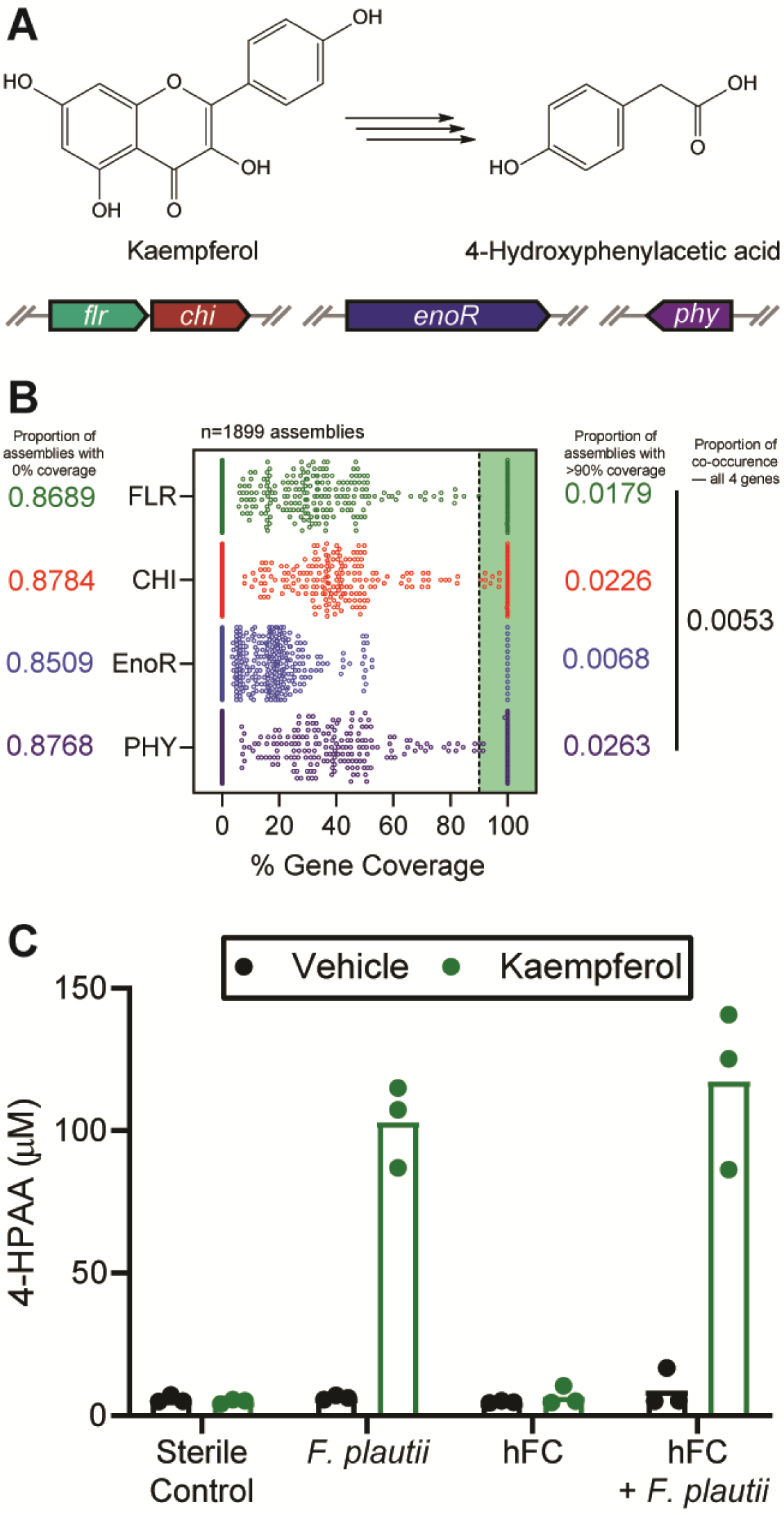
Flavonol and flavone catabolizing genes are rarely found in human microbiomes. (**A**) Graphical depiction of the complete bacterial gene set required to catabolize the flavonol kaempferol into 4-HPAA. (**B**) Computational screening of 1,899 assemblies of metagenomic data derived from human fecal samples representing over 1,300 human subjects for each of the bacterial genes required to catabolize flavonols/flavones. Assemblies with >90% coverage to the parent *Flavonifractor plautii* YL31 gene of interest were considered present. (**C**) *F. plautii* converts kaempferol into 4-HPAA after 24-hours of anerobic incubation in vitro and retains its function when added to a non-converting human fecal community (hFC). Individual points represent individual technical replicates, and bars represent group means.

Having deduced the predicted rarity of flavonol catabolizing genes in human metagenomes, we next set out to test *in vitro* conversion of the flavonol kaempferol into 4-HPAA by stable human-derived polymicrobial communities. In addition, we tested whether community converter capacity could be induced by supplementation with an isolate of *F. plautii* with known flavonol catabolizing capacity (Fig. 5C). In this experiment, we used a human-derived fecal microbiome that lacked catabolic activity toward kaempferol as a negative control and scaffold community into which we spiked in *F. plautii*. Following a 24-hour growth period in anaerobic conditions at 37° C, we provided either a vehicle control or kaempferol substrate and incubated the communities for an additional 24 hours. Using a targeted LC-MS/MS approach to measure 4-HPAA, the complete conversion of kaempferol substrate to 4-HPAA was observed in the *F. plautii* containing community but not the scaffold negative control community. Together, these data support our hypothesis that a single microbe with catabolic activity towards flavonoids can be introduced into complex human fecal microbial communities to produce 4-HPAA *in vitro*, creating perspective for use of *F. plautii* in probiotic interventions.

## Discussion

Dietary intervention is a common approach to combat cardiometabolic disease (CMD), often complementing traditional pharmaceutical therapies. The human gut microbiota extensively metabolizes dietary input, yielding a plethora of microbe-derived metabolites with poorly characterized influences on host physiology. Pertinent to this study, dietary flavonoids have been reported to abrogate diet-induced obesity in a microbe-dependent manner (*8, 9*). Moreover, several commensal gut bacteria are known to catabolize flavonoids into monophenolic acids (*15*). However, the contribution of these diet-derived gut microbial metabolites to CMD remains unclear. In an effort to understand the role of microbial flavonoid catabolites in obesity-related CMD, we used a multifaceted approach to study the interface of diet, the gut microbiota, and host physiology with single-molecule resolution.

Our study uncovers the functional importance of a single flavonoid-derived microbial catabolite, 4-HPAA, in abrogating HFD-induced hepatic steatosis. In addition, we establish the ability of 4-HPAA to activate AMPKα and modulate its downstream effectors. Remarkably, our investigation of the presence of *F. plautii* flavone/flavonol catabolizing (4-HPAA producing) genes present in human metagenomic data revealed that although 28 percent of the 1,899 assemblies contained any one particular strain of *F. plautii*, less than one half of one percent contained all four genes required to produce 4-HPAA from a flavonoid precursor. Paired with the notable intra-individual variation of flavonoid catabolic activity in humans (*29, 30*), these data underscore the need to consider the microbial contribution to dietary intervention as a complementary strategy to combat CMD. Stratification by “responder” status based on the co-occurrence of the complete flavonoid catabolizing gene set may inform, in part, the efficacy of intervention. While theoretically possible, the ability of a probiotic flavonoid-catabolizing *F. plautii* strain to stably colonize the gut remains to be determined and may prove challenging due to strain level competition, particularly in the presence of a flavonoid substrate (*34*), and the frequency of non-synonymous mutations over time (*35*). As an alternative, additional flavonoid-catabolizing microbiota members can be isolated and characterized or a next generation synbiotic could be engineered by inserting the flavonoid-catabolizing gene set into a stably colonizing commensal host.

Our study has several limitations. First, studying human-relevant dietary substrates in a mouse model is reductive in that the human and mouse microbiota are not entirely comparable entities (*36*). Consequently, the development of humanized gnotobiotic mouse models have served as a welcomed advance in the study of diet-gut microbiota interactions (*37*). However, even this state-of-the-art approach has its own limitations as the colonization of the complete donor community is often not possible due to specific microbe-host dependencies (*37*). Second, since germ-free mice are resistant to diet-induced obesity (*38*), we were unable to test the anti-obesogenic properties of elderberry extract-supplemented HFD or 4-HPAA in the absence of the gut microbiota. Third, the possibility exists that undiscovered bacterial catabolic pathways may catabolize flavones/flavonols into 4-HPAA. To this point, we simply used the knowledge at our disposal, focusing on the known catabolic pathway. Lastly, although the subcutaneous administration provides a high level of resolution, this approach discounts the physiological circulation of gut microbial metabolites, first draining through the mesentery into the portal vein before delivery to the liver. Future studies can overcome this challenge by continuously infusing metabolites of interest directly into the portal vein, mirroring the natural circulation of microbial metabolites in vivo (*18*).

In conclusion, we used an array of *in vitro, in vivo*, and *in silico* analyses to reveal the gut microbial contribution of flavonoid catabolism in the context of overnutrition-induced metabolic disease and identified 4-HPAA, a single microbe-derived metabolite sufficient to abrogate obesity-driven hepatic steatosis. We see this as an example of how a single gut microbial metabolite stemming from the diet can profoundly impact host physiology and is a step towards combined diet-probiotic intervention therapies for cardiometabolic diseases.

## Acknowledgments

We would like to thank Brandon Stelter from the Cleveland Clinic Center for Medical Art and Photography for creating the graphical abstract.

## Author contributions

Conceptualization: LJO, JMB, JC

Methodology: LJO, AMH, CM, PPA, JMB, JC

Investigation: LJO, KS, WM, BD, IC, VV, KF, AJH, DO, IN, DSA, NS

Visualization: LJO, JC

Funding acquisition: LJO, LEN, ZW, DSA, AMH, SLH, JMB, JC

Writing – original draft: LJO

Writing – review & editing: All authors

