## Supplemental information for "A gut microbial metabolite of dietary polyphenols reverses obesity-driven hepatic steatosis"

**This PDF file includes:**

Materials and Methods
Figs. S1 to S5

### Materials and Methods

Animal studies

Male C57BL/6 mice were purchased from Jackson Laboratory and housed under specific pathogen-free conditions within the Biological Resources Unit of Cleveland Clinic Lerner Research Institute. For the berry extract feeding study, 6-week-old mice were randomly assigned ad libitum access to a standard high fat diet (Diet D12492, Research Diets, Inc.), or the same diet supplemented with 1% w/w elderberry extract (Product Code 70120034, Artemis International), black currant extract (Product Code 24140500, Artemis International), or aroniaberry extract (Product Code 24198000, Artemis International) for 16 weeks. For the subcutaneous implant experiment, 4-week-old mice were fed a standard high fat diet with 60 kcal% lard-derived fat (Diet D12492, Research Diets, Inc.) ad libitum for 12 weeks to induce obesity-related metabolic syndrome. Next, scaffold control pellets or pellets releasing 350 μg 4-HPAA per day for two weeks were subcutaneously implanted (Innovative Research of America). New pellets were implanted every 14 days for a total of 6 weeks, while the animals remained on ad libitum high fat diet. All experiments and procedures were approved by an Institutional Animal Care and Use Committee.

Phenotyping of mouse models

Body weight and food consumption were measured at baseline and weekly thereafter for the duration of the berry extract feeding experiment and subcutaneous pellet experiment. Lean and fat mass values were assessed at baseline, twenty-five days after the first pellet and at the end of both mouse experiments using EchoMRI body composition analysis (EchoMRI, LLC). Plasma insulin levels were measured using an enzyme-linked immunosorbent assay (ELISA) kit (Product No. 90080, Crystal Chem). Enzymatic analysis of hepatic triglycerides was performed as previously described (*39*). AST and ALT were measured as previously described (*18*). To assess pathologic changes in liver morphology, H&E-stained sections were assessed in a blinded manner by a board-certified pathologist for steatosis, inflammation, and ballooning.

Oxymax/CLAMS metabolic cage studies

For indirect calorimetry studies and activity monitoring, mice (n=6 per group) were single-housed in the Oxymax/Comprehensive Animal Monitoring System (CLAMS) metabolic cage monitoring system (Columbus Instruments) 25 days after the initial pellet implant. The animals were allowed to acclimate for 24 hours under murine isothermal conditions (*40*) (30° C) before 24 hours of continuous data monitoring at 30° C, 24 hours at 23° C, and 24 hours at 4° C. Data analysis was performed as previously described(*18*) using CalR (*41*).

Cecal microbiome sequencing and analysis

DNA was extracted from mouse cecal contents and food samples using the QIAGEN PowerSoil Pro kit using the manufacturer’s protocol. 16S rRNA amplicon sequencing was done for the V3-V4 region using Illumina iSeq 100 system from mouse cecal contents. Raw 16S amplicon sequences and metadata were demultiplexed using the split_libraries_fastq.py script implemented in QIIME1.9.1(*42*). The demultiplexed fastq file was split into sample specific fastq files using the split_sequence_file_on_sample_ids.py script from QIIME1.9.1 (*42*). Individual fastq files without non-biological nucleotides were processed using the Divisive Amplicon Denoising Algorithm (DADA) pipeline (*43*). The output of the DADA2 pipeline (feature table of amplicon sequence variants) was processed for alpha and beta diversity analysis using the phyloseq (*44*) and microbiomeSeq (http://www.github.com/umerijaz/microbiomeSeq) packages in R. Alpha diversity estimates were measured within group categories using the estimate_richness function of the phyloseq package (*44*). Non-metric multidimensional scaling (NMDS) was performed using the Bray-Curtis dissimilarity matrix (*45*) between groups and visualized by using the ggplot2 package (*46*). We assessed the statistical significance ($p < 0.05$) throughout, and whenever necessary we adjusted p-values for multiple comparisons according to the Benjamini-Hochberg method to control false discovery rate (FDR) (*47*) while performing multiple testing on taxa abundance according to sample categories. We performed an analysis of variance (ANOVA) among sample categories while measuring the alpha diversity measures using the plot_anova_diversity function in the microbiomeSeq package (http://www.github.com/umerijaz/microbiomeSeq). Permutational multivariate analysis of variance (PERMANOVA) with 999 permutations was performed on all principal coordinates obtained during principal coordinates analysis with the ordination function of the microbiomeSeq package. Linear regression (parametric) and Wilcoxon (non-parametric) tests were performed on amplicon sequence variant abundances against meta-data variable levels using their base functions in R (*48*).

Human microbiome metagenomic analysis

Longitudinal shotgun microbiome data from Poyet et. Al., 2019 (*32*) and Zou et al., 2019 (*33*) was downloaded using Entrezpy[44]. Shotgun microbiome data was quality trimmed using nesoni pipeline (https://github.com/Victorian-Bioinformatics-Consortium/nesoni). MetaPhlAn2 (METAgenomic PHyLogenetic ANalysis) was used for species level metagenomic profiling of the quality trimmed reads (*50*). Using megahit assembler shotgun microbiome data was assembled into contigs (*51*). Contigs were screened for flavone reductase (EC 1.1.1.234), chalcone isomerase (EC 5.5.1.6), enoate reductase (EC 1.3.1.3) and phloretin hydrolase (EC 3.7.1.4) using abricate (https://github.com/tseemann/abricate). Quality trimmed reads were also mapped on the nucleotide sequences of flavone reductase (EC 1.1.1.234), chalcone isomerase (EC 5.5.1.6), enoate reductase (EC 1.3.1.3) and phloretin hydrolase (EC 3.7.1.4) using BBMap (https://jgi.doe.gov/data-and-tools/bbtools/bb-tools-user-guide/bbmap-guide/). Count matrix was generated using featurecount function implemented in Subread package (*52*, *53*).

LC-MS/MS analysis of monophenolic acids

Plasma, liver, and bacterial supernatant were prepared for LC-MS/MS analysis and quantified as previously described (*18*). Briefly, stable isotope dilution high performance liquid chromatography tandem mass spectrometry (LC–MS/MS) was used for quantification of levels of 4-HPAA using an AB SCIEX Q-Trap 4000 triple quadrupole mass spectrometer equipped with an electrospray ionization source operating in negative ion mode. Bacterial supernatants were prepared by first pelleting 24-hour cultures by spinning at 15,000 x g for 10 minutes. Next, up to 500 µL was transferred to an Amicon Ultra-0.5 mL 3K filter (Millipore Catalog No. UFC500324) and centrifuged at 15,000 x g for 30 minutes. 20 µL of the filtered supernatant was prepared as previously described for analysis (*18*). The d6(methyl)- isotopologue of 4-HPAA was used as an internal standard (CDN Isotopes Product No. D-7842) at 10 µM for plasma and liver samples and 50 µM for bacterial supernatant. 4-HPAA was monitored using multiple reaction monitoring of precursor and characteristic product ions in negative mode as follows: m/z 151.2 → 107.0 for 4-HPAA; m/z 157.2 → 113.0 for d6-4-HPAA; m/z 120.7 → 77.0 for benzoic acid; m/z 136.8 → 92.9 for 3-hydroxybenzoic acid (3-HBA); m/z 136.8 → 92.9 for 4-

hydroxybenzoic acid (4-HBA); m/z 148.9 → 104.9 for 3-PPA 3-phenylpropanoic acid (3-HPPA); m/z 166.6 → 151.9 for 4-hydroxy-3-methoxybenzoic acid (4-H-3-MBA).

Cell culture and immunoblotting

Primary hepatocytes were isolated from two male C57BL/6 mice using methods described elsewhere (*54*) and plated in 6-well collagen I coated plates at a density of 3 x $10^5$ cells/well in Williams' E medium containing 10% fetal bovine serum. Primary hepatocytes were serum-starved in plain William's E medium for 2-hours prior to treatment with a DMSO vehicle control or 4-HPAA (TCI America Product No. H0290) for 30 minutes. Cell pellets or whole liver homogenates prepared from primary hepatocytes or murine liver, respectively, in a modified RIPA buffer with protease and phosphatase inhibitors. Western blotting was performed as previously described (*55*). Images were captured using a GE Amhersham Imager 6000 and the accompanying software (version 1,1.1) was used for densitometric protein expression analysis. Primary antibodies (pS79-ACC, Cell Signaling, #11818; ACC, Cell Signaling, #3676; pT172-AMPKα, Cell Signaling, # 2535; AMPKα, Cell Signaling, #5831; pS182-AMPKβ1, Cell Signaling, #4186; AMPKβ1, Cell Signaling, #4178) were prepared 1:1000 in TBST buffer with 5% (w/v) BSA. Rabbit secondary (Cell Signaling, #7074) and β-actin (Cell Signaling, #12620) antibodies were prepared 1:5000 in TBST with 5% (w/v) non-fat dried milk.

Microbial culturing

Microbial culture experiments were performed in an anaerobic chamber (Coy Laboratory Products, Inc.) under the following conditions: 85% $N_2$, 5% $H_2$,10% $CO_2$, and < 25 PPM $O_2$. To examine the conversion of kaempferol substrate into 4-HPAA, a roughly 1 cm section of human fecal material prepared using the FAST technique (*56*) was suspended in 10 mL of a 50:50 (w:w) mixture of Wilkins-Chalgren (*57*) and Gifu Anaerobic Medium (*58*), vortexed for 2 minutes, and incubated at 37° C for 24 hours. Next, the resulting microbial culture was mixed 1:1 (v:v) with fresh media containing either DMSO vehicle or kaempferol hydrate (TCI America Product No. K0018) at a final concentration of 100 μM and cultured at 37° C in a deep-well 96-well plate for 24 hours. The reaction was stopped by freezing the 96-well plate at -80° C.

Whole genome sequencing and analysis

DNA was extracted from *Flavonifractor plautii* ATCC 49531 using the QIAGEN DNeasy PowerSoil Pro kit. Illumina library preparation preceded 2 x 151 bp sequencing using the NextSeq 2000 platform. The reads were trimmed and quality controlled using Trimmomatic (*59*) and then assembled using SPAdes (*60*) and annotated via Prokka (*61*).Genome visualization, annotation, and phylogenetic comparisons were using PATRIC with RASTk (*62, 63*). The closest reference and representative genomes were identified by Mash/MinHash (*64*). PATRIC global protein families (PGFams) (*65*) were selected from these genomes to determine the phylogenetic placement of this genome. The protein sequences from these families were aligned with MUSCLE (*66*), and the nucleotides for each of those sequences were mapped to the protein alignment. The joint set of amino acid and nucleotide alignments were concatenated into a data matrix, and RaxML (*67*) was used to analyze this matrix, with fast bootstrapping was used to generate the support values in the tree (*68*).

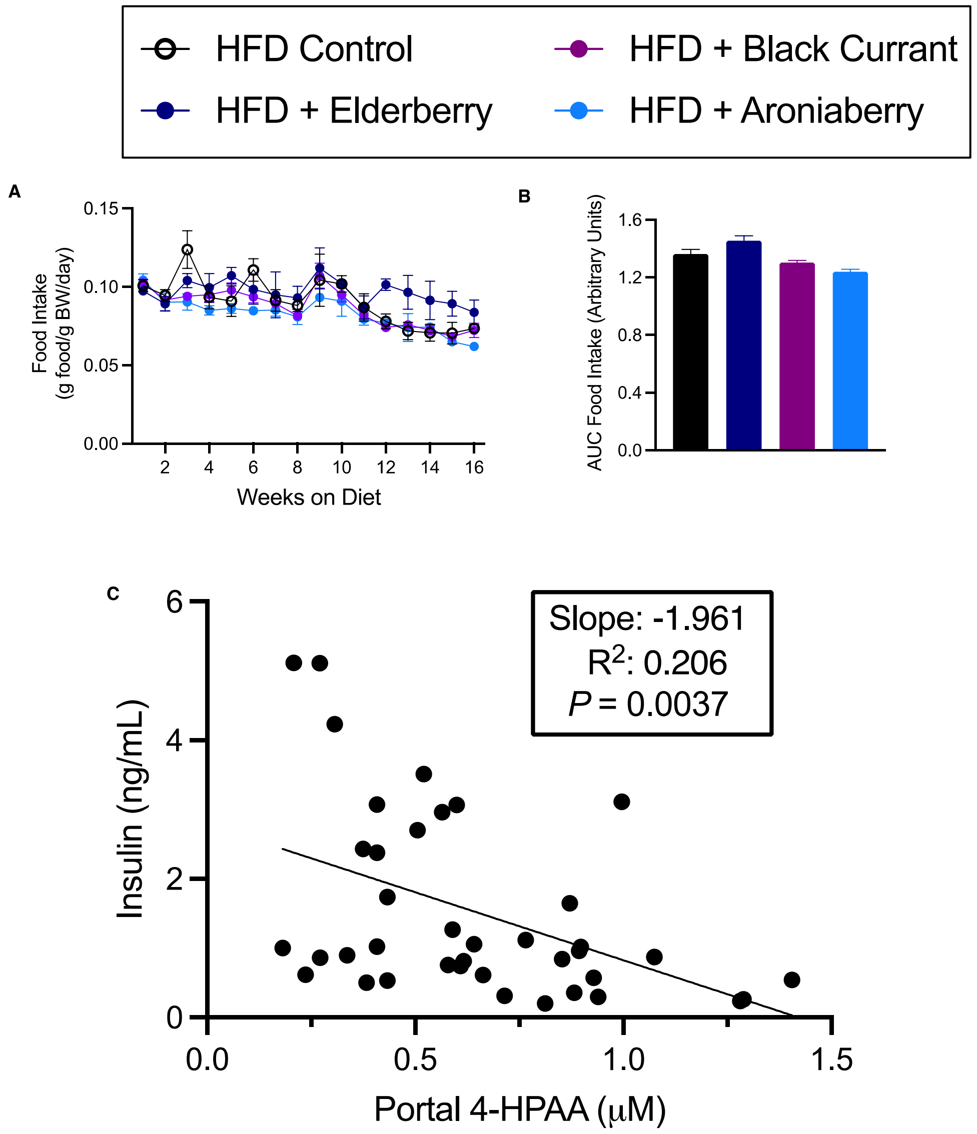


**Fig. S1. Berry extract supplementation does not impact food consumption but modulates the gut microbial community structure.** (**A**) Weekly food consumption data represented as grams of food consumed per gram of body weight per day. (**B**) Mean cumulative AUC for food intake as represented in panel A; ordinary one-way ANOVA with Dunnett's multiple comparisons test with error bars representing SEM. n=9-10 per group. (**C**) Simple linear regression analysis of terminal plasma insulin and portal plasma 4-HPAA. Individual points represent individual mice, and bars represent group means.

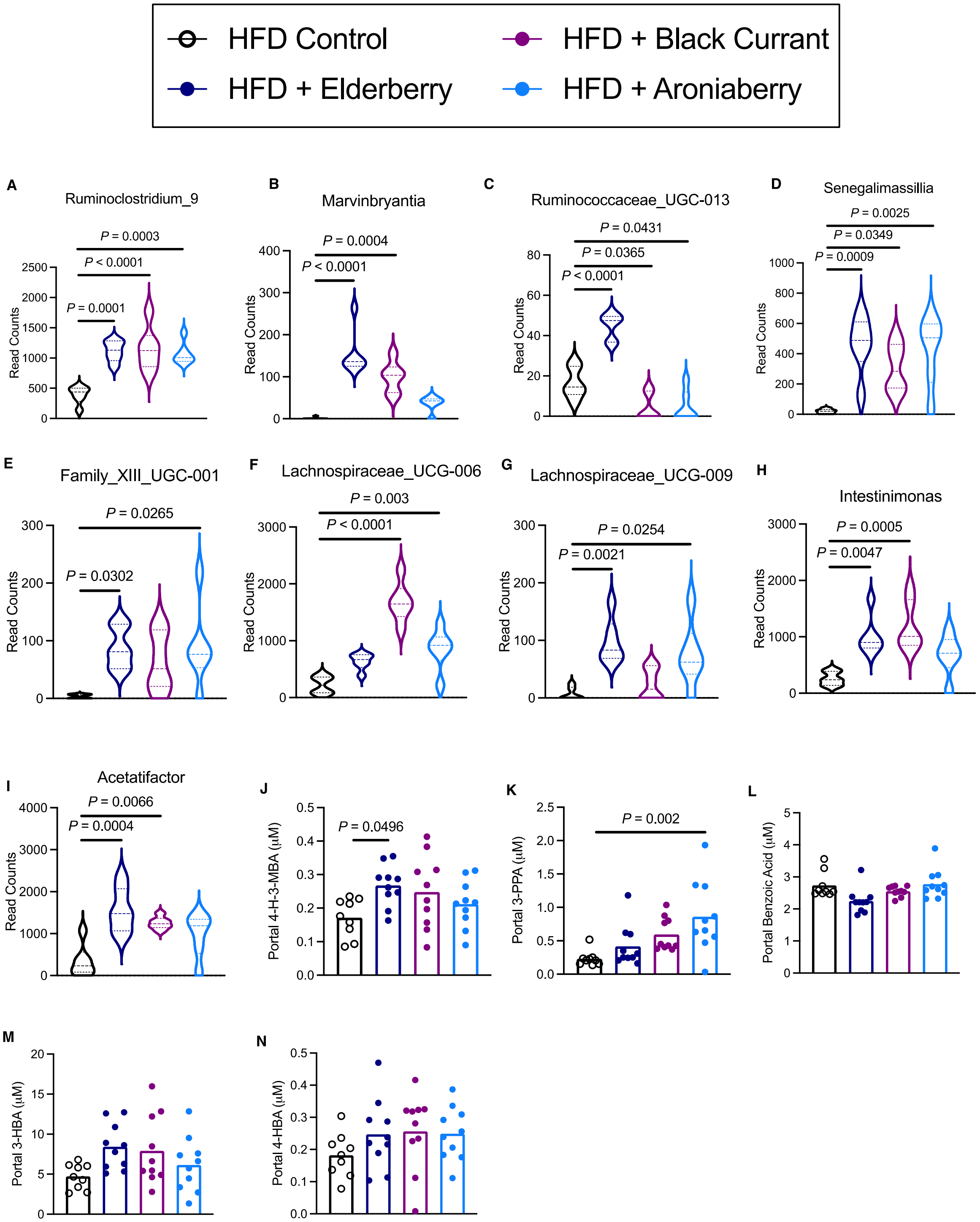


**Fig. S2. Berry extract feeding impacts cecal microbiome and portal plasma flavonoid catabolite levels.** (**A-I**) Violin plots of representative differentially abundant bacterial genera identified via 16S rRNA sequencing of cecal contents at the time of necropsy, n=6 per group. Ordinary one-way ANOVA with Tukey's multiple comparisons test used to analyze**.** (**J-N**) Portal plasma concentration of microbial flavonol catabolites measured by LC-MS/MS, one-way ANOVA with Dunnett's multiple comparisons test; n=9-10 per group. Individual points represent individual mice, and bars represent group means.

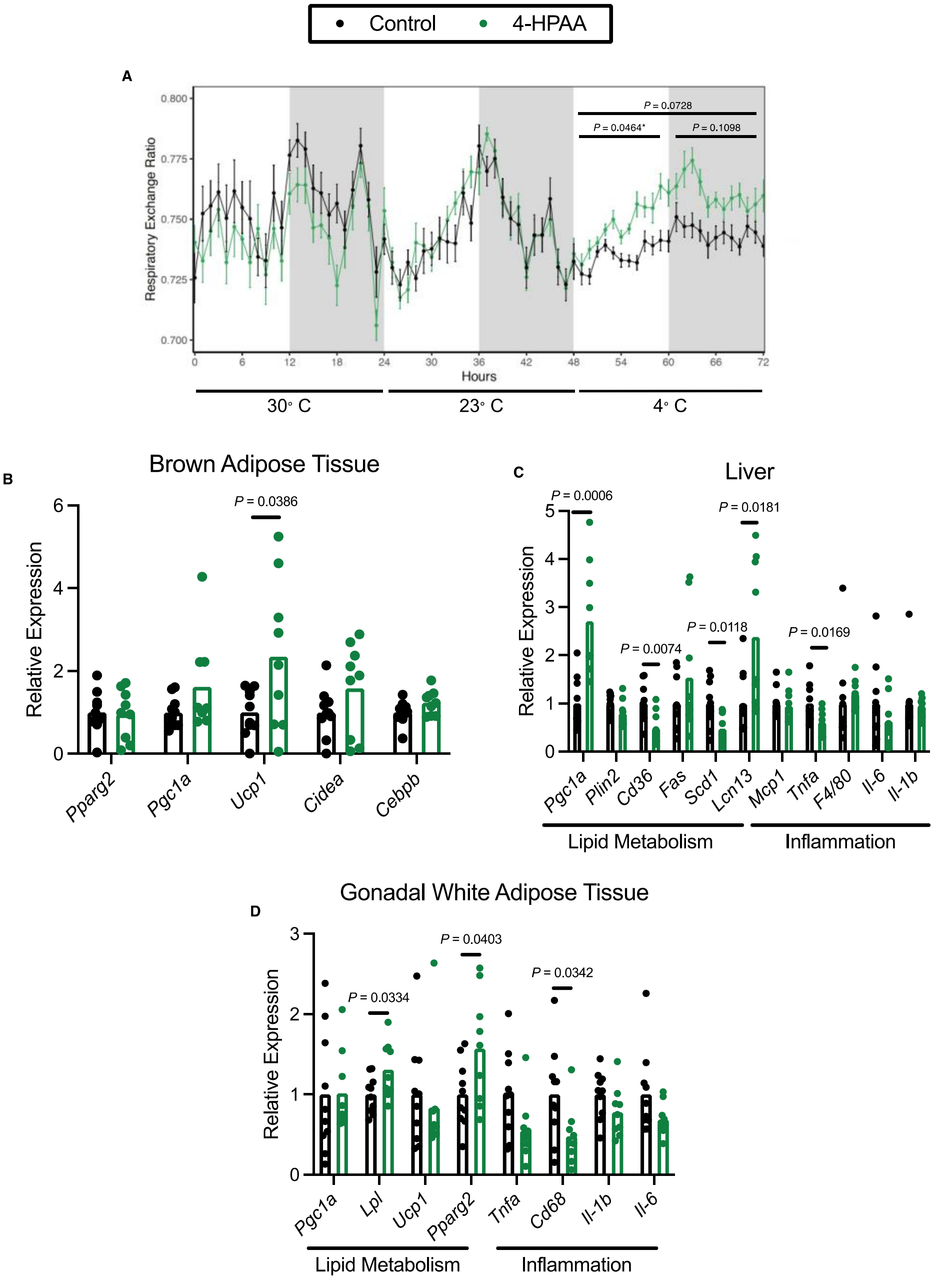


**Fig. S3. 4-HPAA treated mice demonstrate improved cold tolerance and transcriptional changes in metabolically active peripheral tissues.** (**A**) After twenty-five days of 4-HPAA treatment, global energy substrate utilization was assessed using the Oxymax/CLAMS metabolic cages system, n=6 per group. (**B-D**) RT-qPCR quantification of mRNA transcripts involved in global metabolic programming and inflammation in brown adipose tissue normalized to the average of *Actb* and *Hprt* (**B**), the liver normalized to *Ppia* (**C**), and gonadal white adipose tissue normalized to *Tbp or Ppia* (**D**), n=9-10 per group. Statistical analysis of Oxymax/CLAMS data was performed using ANOVA in CalR (***41***). Statistical analysis of RT-qPCR data was performed using unpaired two-tailed Student’s t-test. Individual points represent individual mice, and bars represent group means.

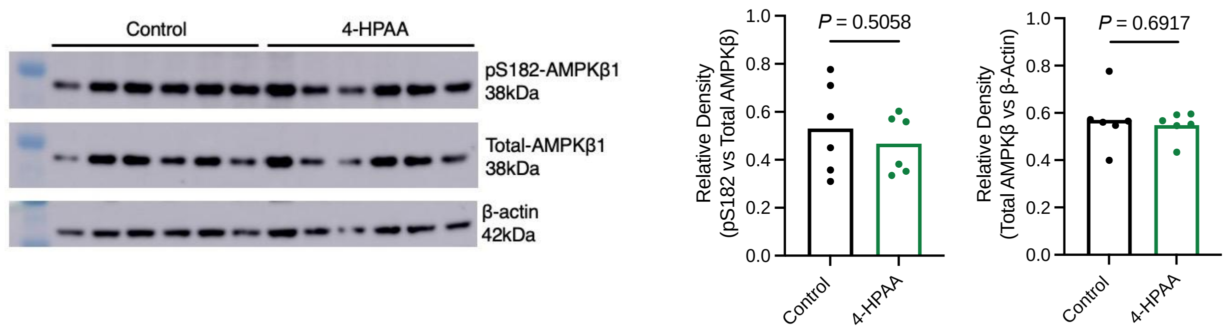


**Fig. S4. 4-HPAA does not activate AMPKb in mice.** Western blot analysis of pAMPKβ1 (S182), total AMPKβ1, and β-actin with densitometric quantification, n=6 per group. Statistical analysis was performed using an unpaired two-tailed Student's t test. Each dot represents an individual mouse.

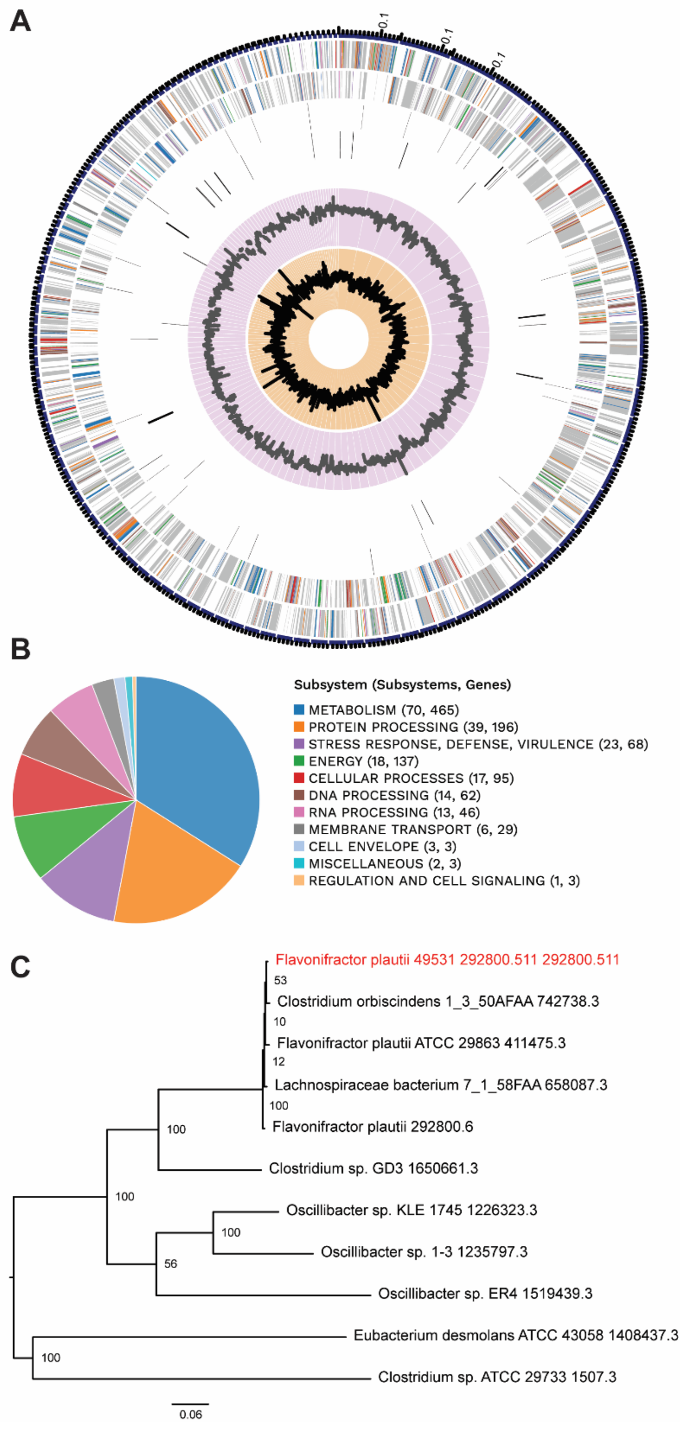


**Fig. S5. *Flavonifractor plautii* ATCC 49531 genome composition and phylogeny.** (**A**) A circular graphical display of the distribution of the genome annotations. From outer to inner rings, the contigs, CDS on the forward strand, CDS on the reverse strand, RNA genes, CDS with homology to known antimicrobial resistance genes, CDS with homology to know virulence factors, GC content and GC skew. The colors of the CDS on the forward and reverse strand indicate the subsystem that these genes belong to as summarized in (**B**). Only the largest 129 contigs are shown. (**C**) Phylogenetic analysis of the *F. plautii* ATCC 49531 genome.
